# Plant-based Production and Characterization of a Promising Fc-fusion Protein against Bone Mass Density Loss

**DOI:** 10.1101/2022.06.05.494914

**Authors:** Yongao Xiong, Hiroto Hirano, Nancy E. Lane, Somen Nandi, Karen A. McDonald

**Author notes:** **Correspondence:** Karen McDonald.

## Abstract

Microgravity-induced bone loss is a main obstacle for long term space missions as it is difficult to maintain bone mass when loading stimuli is reduced. With a typical bone mineral density loss of 1.5% per month of microgravity exposure, the chances for osteoporosis and fractures may endanger astronauts’ health. Parathyroid Hormone or PTH (1-34) is an FDA approved treatment for osteoporosis, and may reverse microgravity-induced bone loss. However, PTH proteins requires refrigeration, daily subcutaneous injection, and have a short shelf-life, limiting its use in a resource-limited environment, like space. In this study, PTH was produced in an Fc-fusion form via transient expression in plants, to improve the circulatory half-life which reduces dosing frequency and to simplify purification if needed. Plant-based expression is well-suited for space medicine application given its low resource consumption and short expression timeline. The PTH-Fc accumulation profile in plant was established with a peak expression on day 5 post infiltration of 373 ± 59 mg/kg leaf fresh weight. Once the PTH-Fc was purified, the amino acid sequence and the binding affinity to its target, PTH 1 receptor (PTH1R), was determined utilizing biolayer interferometry (BLI). The binding affinity between PTH-Fc and PTH1R was 2.30 × 10^−6^ M, similar to the affinity between PTH (1–34) and PTH1R (2.31 × 10^−6^ M). Its function was also confirmed in a cell-based receptor stimulation assay, where PTH-Fc was able to stimulate the PTH1R producing cyclic adenosine monophosphate (cAMP) with an EC_50_ of (8.54 ± 0.12) × 10^−9^ M, comparable to the EC_50_ from the PTH (1-34) of 1.49 × 10^−8^ M. These results suggest that plant recombinant PTH-Fc exhibits a similar potency compared to PTH. Furthermore, it can be produced rapidly at high levels with minimal resources and reagents, making it ideal for production in low resource environments such as space.

## 1 Introduction

Microgravity-induced bone loss is a main obstacle for long term space missions. Bone is a dynamic organ that serves important mechanical and calcium homeostatic functions (Demontiero et al., 2012). It constantly undergoes a self-regeneration process called bone remodeling, a process in which the old bone is resorbed by osteoclasts and new bone is regenerated by osteoblasts. Maintaining a balance between bone resorption and regeneration is critical for human health. For astronauts exposed to microgravity, it is very difficult to maintain such a balance; as the loading stimuli reduces, there is an increase in bone resorption with no change or even decreased bone formation, leading to bone density loss (Ohshima, 2012). Microgravity-induced bone loss became a concern since Gemini flights (1-14 day durations, 1962 – 1966), where “small but significant” bone loss was reported with less than two weeks of microgravity exposure (MACK and LaChance, 1967; Mack et al., 1967). Although the percentage of bone mineral density (BMD) loss was measured using densitometry of plain X-rays in these studies, which was not an accurate methodology, it raised the awareness of detrimental effects of microgravity on bone density. As technology advanced, the typically BMD loss was determined to be 1.5% per month of microgravity exposure (Ohshima, 2012), which endangers astronauts’ health with an increased chance for osteoporosis and fractures, especially during extravehicular activities. Physical exercise as a countermeasure for BMD loss is effective but cannot eliminate the problem completely (Ohshima, 2012). Bone regenerative therapeutics such as PTH (1-34, Forteo^®^), are FDA approved treatments for osteoporosis, and have vast potential to reverse microgravity-induced bone loss. PTH stimulates bone regeneration by activating osteoblast cells through binding to the PTH1R on its cell surface, and promotes bone formation. However, PTH requires refrigeration, daily subcutaneous injection, and has a short shelf-life, limiting its use in a resource-limited environment, like space. To improve circulatory half-life of PTH and minimize resources utilized for protein production in space, PTH was expressed in a Fc-fusion form, namely PTH-Fc in plants transiently. The addition of the Fc component will likely allow for a longer circulatory half-life by interaction with the salvage neonatal Fc-receptor (Roopenian and Akilesh, 2007) and avoids frequent redosing. Studies using a recombinant PTH-Fc fusion protein produced in *E. coli* showed a 33-fold increase in mean circulation residence time in rats as well as significant increases in bone volume, density and strength in osteopenic mice and rats (Kostenuik et al., 2007). A single dose of a fusion between PTH (1–33) and the collagen binding domain (PTH-CBD), designed to increase circulatory half-life, sustained increases in BMD by >10% in normal mice for up to a year (Ponnapakkam et al., 2012).

Unlike most other biologic production platforms, producing biologics in plants transiently requires only growing plants and an expression vector to deliver the gene. In this study, *Agrobacterium tumefaciens* (*Agrobacterium*) was used to deliver a PTH-Fc expressing cassette into plant tissue by vacuum infiltration given its horizontal gene transfer capacity. Alternatively, a plant virus (Kawakami et al., 2004) or particle bombardment (Kikkert et al., 2005) can be used for gene delivery to further reduce resource requirements. In addition to its low resource requirement, plants present no risk of mammalian pathogen infection and are capable of post-translational modifications, making it a well-suited platform for biologics production under resource constraints.

PTH-Fc forms a homodimer under physiological conditions via disulfide bridges within the Fc hinge region. The Fc domain serves solely as a serum half-life enhancer in this fusion protein; thus, the Fc N-glycosylation site was removed to avoid potential Fc effector functions and associated inflammatory responses. In addition, a SEKDEL C-terminal motif was included to target PTH-Fc for ER retention, which often results in a higher protein yield than targeting proteins for secretion (Pan et al., 2008; Sainsbury and Lomonossoff, 2008). In this study, the PTH-Fc expression, integrity, receptor binding affinity and receptor stimulation efficacy were evaluated, and the biological activity of PTH-Fc was compared to the PTH (1 – 34) peptide alone. To our knowledge, this is the first report of recombinant protein of PTH-Fc in plants.

## 2 Materials and Methods

### 2.1 PTH-Fc Construct

PTH-Fc was expressed in a replicating binary vector as shown in Figure 1. The PTH-Fc fusion protein sequence consists of PTH (amino acids 1 – 34, UniProtKB: P01270), a flexible linker (GGGGS), the Fc region of human IgG1 (amino acid 108 – 329, Genbank: AAC82527.1) with a point mutation (N111Q on PTH-Fc) making the protein aglycosylated, followed by a C-terminal ER retention motif of SEKDEL. The PTH-Fc coding sequence was cloned into the replicating vector with gemini viral components (LIR, SIR, C1 and C2) allowing for replication of the DNA fragment flanked by the LIR domains by rolling circle replication process upon agroinfiltration (Chen et al., 2011). The resulting PTH-Fc expressing binary vector was used to transform *Agrobacterium tumefaciens* EHA105 with the helper plasmid (pCH32) via the freeze-thaw method, resulting in a PTH-Fc expressing *Agrobacterium* strain, pRI201_Gemini_PTH-Fc.

**Figure 1.**
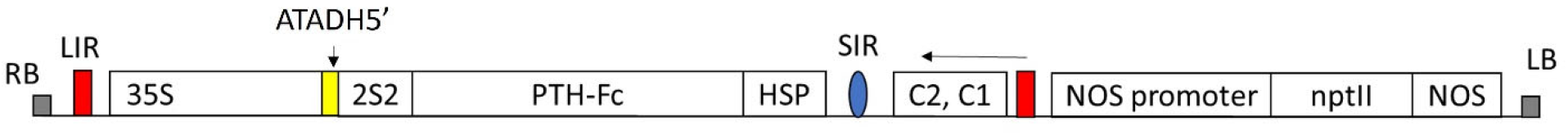
The gene construct for transient PTH-Fc expression in *Nicotiana benthamiana*. RB/LB: right border/left border; LIR: long intergenic region; 35S: Cauliflower mosaic virus 35S promoter; ATADH5’: 5’ UTR of the *Arabidopsis thaliana* alcohol dehydrogenase gene for translation enhancement; 2S2: secretory signal peptide; PTH-Fc: PTH-Fc coding sequence; HSP: *Arabidopsis thaliana* HSP 18.2 terminator; SIR: short intergenic region; C1,C2: Rep/RepA coding sequence; NOS promoter/NOS: nopaline synthase promoter/terminator. nptII: gene codes for an aminoglycoside phosphotransferase II conferring resistance to kanamycin for stable transgenic line selection (if desired).

### 2.2 Transient PTH-Fc Production in *Nicotiana benthamiana*

Transient PTH-Fc production was performed as described previously (Xiong et al., 2019). In brief, *Agrobacterium* strains containing the PTH-Fc expression cassette (pRI201_Gemini_PTH-Fc) and RNA gene silencing suppressor P19 were cultured separately in LB media with appropriate antibiotics and were suspended into the infiltration buffer (10 mM MES buffer at pH 5.6, 10 mM MgCl_2_ and 150 μM acetosyringone, and 0.02% v/v Silwet-L-77) with a final cell density of 0.25 (O.D.600) for each strain. *Nicotiana benthamiana* plants (6-7 weeks old) were vacuum infiltrated with the *Agrobacterium* suspension for 2 min, followed by plant incubation for up to 6 days allowing for protein accumulation.

### 2.3 Protein Extraction and Purification

Harvested plant tissue stored at -80 °C was ground to fine powder with liquid nitrogen using mortar and pestle, and the extraction buffer (PBS, pH 7.4 with 1mM EDTA and 2mM sodium metabisulfite) was added to the leaf powder at a leaf mass (g) to buffer volume (mL) ratio of 1:4. The mixture was incubated at 4°C for 30 min with shaking, filtered through cheesecloth and centrifuged at 35,000 x g for 20 min, followed by 0.22 µm filtration. The filtered crude plant extract was loaded onto a Protein A affinity chromatography column, and PTH-Fc was eluted with 100 mM glycine-HCl at pH of 3.0. Purified PTH-Fc was titrated to neutral pH with 1M Tris, pH 11 and dialyzed against PBS overnight at 4°C, followed by storage at -80 °C.

### 2.4 ELISA Quantification Of PTH-Fc in Crude Plant Extract

PTH-Fc in crude extract was quantified using a direct ELISA. The crude plant extract samples and serial diluted CMG2-Fc standards (from 7.8 µg/mL, 3X serial dilutions) were loaded to a 96-well ELISA microplate and incubated for 1 hr at room temperature (RT), and then blocked with 1% casein in PBS for 30 min (150 µL/well). The bound PTH-Fc was detected with a goat anti-human IgG-HRP antibody at 0.5 µg/mL for 1hr at RT, followed by color development with TMB substrate for 10 min at RT and addition of 1N HCl to stop the reaction. Between steps, prior to color development, plates were washed with 250 µL of PBST 3 times for 5 min each. All the incubation steps were done with 100 µL per well unless otherwise noted. The absorbance at 450 nm was measured with a Spectramax M2 plate reader (Molecular Devices, San Jose, CA) for PTH-Fc quantification.

### 2.5 SDS-PAGE and Western Blotting

Crude extract and purified PTH-Fc were subjected to SDS-PAGE and Western blot analyses. Ten µL of 4X Laemmi dye (non-reducing) or 8 µL 4X Laemmi dye with 2 µL of BME (reducing condition) was added to samples diluted in dd H_2_O to a total volume of 40 µL. The mixtures were then heated at 95 °C for 5 min for protein denaturing, and samples were run on 4 – 20% precast stain free polyacrylamide gels at 200 V for 35 min. Gels were imaged with the ChemiDoc imager (Bio-Rad, Hercules, CA). For Western blotting, gels were then transferred to a 0.2 µm nitrocellulose membrane using the Trans-Blot Turbo Transfer System with the “MIXED MW” protocol. The membranes were blocked with 1% casein in PBS for 1 hr at RT, washed 3 times in PBST for 5 min each, and then probed with a mouse anti-PTH antibody (1:1000) or a mouse anti-KDEL antibody (1:1000) for 1 hr at RT. The membranes were washed 3 times for 5 min each before incubating with a goat anti-mouse-HRP antibody (1:2,000) for 1 hr at RT. After 3 washes, membranes were developed with Clarity Western ECL substrate and imaged with the ChemiDoc imager under chemiluminescent blot settings.

### 2.6 Protein sequence identification by Liquid Chromatography Tandem Mass Spectrometry (LC-MS/MS)

Purified PTH-Fc was sent to the UC Davis Proteomics Core facility for protein sequence identification using LC-MS/MS as described previously (Xiong et al., 2019).

### 2.7 Binding affinity analysis by biolayer interferometry

Ni-NTA sensor tips were hydrated in the Kinetics buffer for 10 min and dipped into his-tagged PTH1R diluted in the Kinetics buffer at 1 µM for 1 hr with constant shaking. All steps were performed at room temperature. The functionalized sensor tips were then mounted onto a sensor rack and placed in the Octet 384RED (Fortébio, Fremont, CA) sensor tray. PTH-Fc, PTH 1-34 amino acids (positive control) and CMG2-Fc (negative control) were 2X serial diluted from 10 µM in Kinetics buffer and loaded onto a non-binding black 96-well plate at 200 µL per well. The sensor tips were dipped into analyte solutions for 1,500 s after a 300 s baseline step in the kinetics buffer, and then switched to the kinetics buffer for 1,800 s allowing for dissociation. The sensorgrams were fitted to a 1:1 binding model, and the average responses at the end of the association phase (1,490 – 1,495 s) were used for steady state analysis using ForteBio Data Analysis software.

### 2.8 Receptor stimulation cell-based assay

The cell-based assay (CBA) was performed following manufacturer’s protocol (cAMP Hunter eXpress GPCR Assay). Briefly, CHO-K1 cells expressing PTH1R on the cell surface were seeded onto a black well, clear bottle tissue culture treated 96-well plate at a density of 3.125 × 10^5^ cells/mL with cell plate reagent and incubated at 37 °C, 5% CO_2_ for 24 hr. The cell plating reagent was discarded and replaced with 30 µL of cell assay buffer and then treated with 15 µL 3X serial diluted PTH-Fc (from 9 µM) for 30 min at 37 °C, 5% CO_2_. Anti-cAMP antibody solution (15 µL/well) and 60 µL/well of cAMP working detection solution (contains the enzyme donor) were sequentially added to all wells, and incubated at RT for 1 hr in dark. The cAMP solution A (contains the enzyme acceptor) was then added at 60 µL/well and incubated for 3 hr at RT in dark. The plate was read with a SpectraMax M2 plate reader under luminescent settings, and the EC_50_ was estimated by fitting the dose-response curve to the [Agonist] vs response (three parameters) model in GraphPad Prism 8.

### 3 Results

### 3.1 Transient Expression of PTH-Fc in plants

PTH-Fc was transiently expressed in *Nicotiana benthamiana* whole plants *via* agroinfiltration, and the expression level from 1-6 days post infiltration (DPIs) was determined in crude plant extract with a direct ELISA detecting the Fc domain of PTH-Fc. Protein expression (Figure 2 A) was first detected on 2 DPI and continued to increase until 5 DPI, reaching a maximum expression level of 373 ± 59 mg/kg leaf fresh weight; after 5 DPI the protein level started to drop. This expression level is comparable to another Fc-fusion protein produced transiently in *Nicotiana benthamiana* (Xiong et al., 2019). These results suggest that 5 DPI is the optimal harvesting time for PTH-Fc, and protein degradation in the plant exceeds production with a longer incubation period. The Western blot analysis on crude extracts (Figure 2 B) from 1 to 6 DPI confirmed the presence of the PTH domain, with a band height around the theoretical molecular weight of PTH-Fc monomer at 31.1 kDa. The higher band between 50 and 75 kDa in samples from 3, 4 and 5 DPIs represents the nonreduced PTH-Fc dimer.

**Figure 2.**
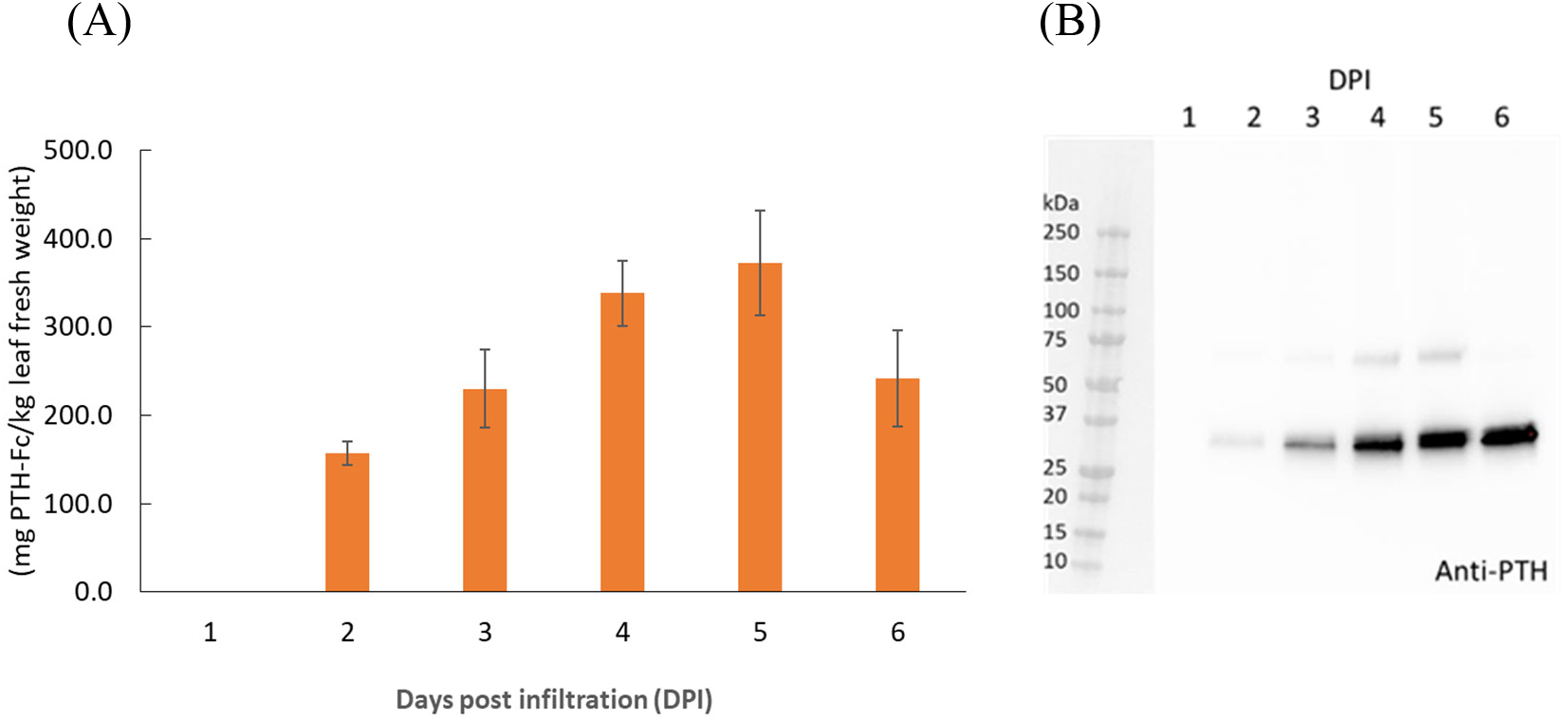
PTH-Fc in plant accumulation profile from 1 – 6 DPI by ELISA (A) and Western blot analysis detecting PTH domain of PTH-Fc in crude plant extracts from 1 – 6 DPI (B). Error bars represent the standard error of the mean of duplicate measurements.

### 3.2 PTH-Fc Amino Acid Sequence Identification by LC-MS/MS

To confirm protein integrity and amino acid sequence, Protein A purified PTH-Fc was subjected to LC-MS/MS analysis. The control sequence (expected amino acid sequence) and detected sequence in PTH-Fc sample are shown in Figure 3 with the sequences not detected represented with dashes. The 2S2 secretory signal peptide is underlined in the control sequence, which is expected to be removed upon protein maturation. The sequence coverage of PTH-Fc with respect to the control sequence was 75.7% (2S2 excluded). Signal peptide was not detected in the PTH-Fc sample, indicating that it was correctly removed from the mature PTH-Fc. However, a 50 amino acid portion at the C-terminus was not detected in LC-MS/MS, thus, Western blot analysis detecting the C-terminal motif, SEKDEL, was performed to confirm protein integrity.

**Figure 3.**
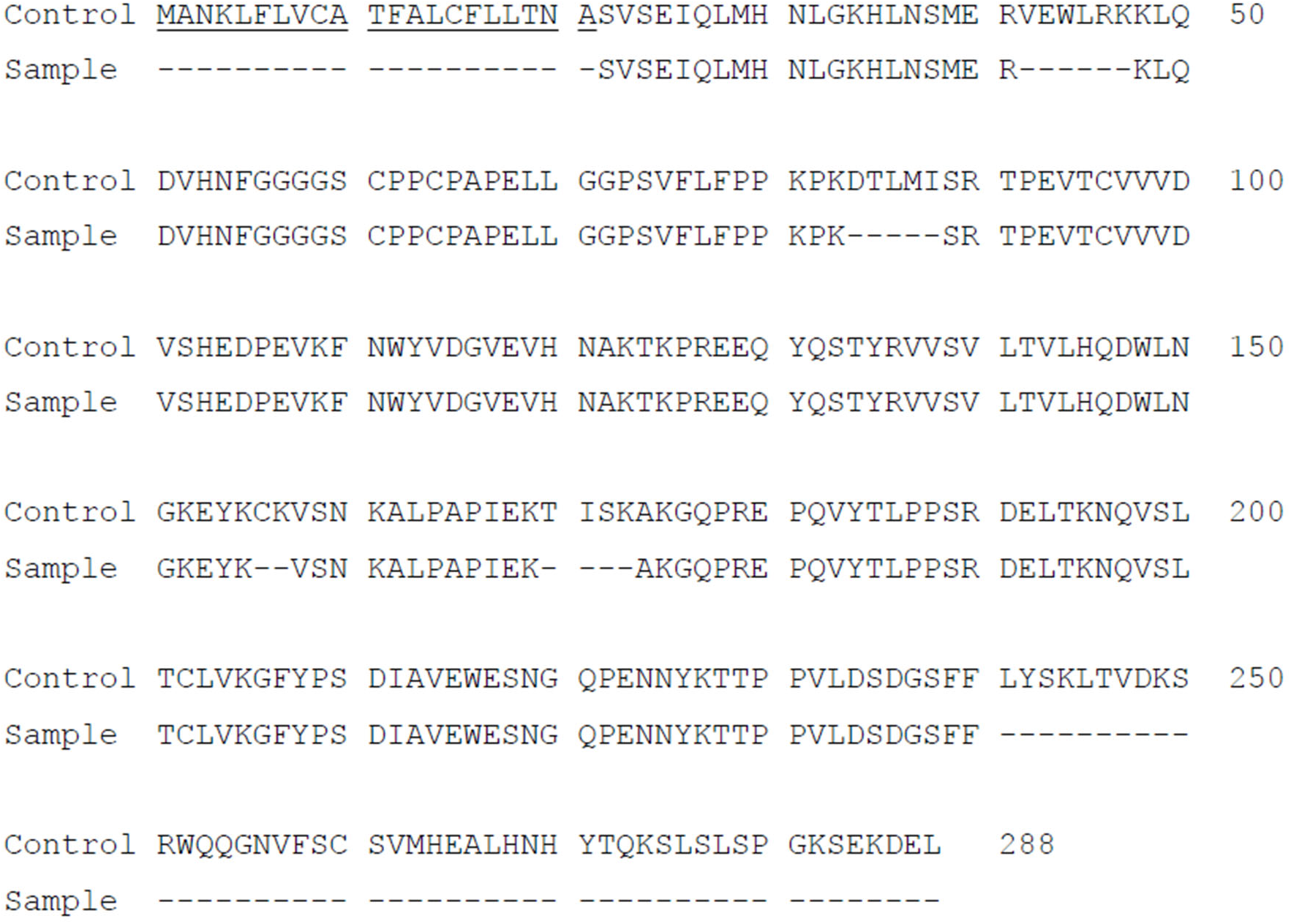
Amino acid sequence of PTH-Fc compared to the control sequence with undetected sequences represented as dashes. The underlined sequence corresponds to 2S2 secretory signal peptide.

### 3.3 Western Blotting and SDS-PAGE Analyses on Purified PTH-Fc

As shown in Figure 6.3, PTH-Fc (PTH: 4.1 kDa; linker +Fc: 27 kDa, PTH-Fc: 31.1 kDa) and a positive control protein (47.2 kDa) containing a C-terminal SEKDEL motif were probed by an anti-PTH antibody and an anti-SEKDEL antibody. On the anti-PTH Western blot (Figure 4 A), only the PTH-Fc sample shows a band between 25 and 37 kDa. The faint band in the control lane was due to the PTH-Fc sample overflowed from the adjacent lane. On the anti-SEKDEL Western blot (Figure 4 B), intense bands in both lanes at their expected molecular weights are present, which confirms the presence of SEKDEL sequence in both samples. The missing coverage in MS analysis can be a result of low enzyme efficiency against the C-terminal sequence or low column yield of those peptides that are too hydrophilic or small, which passed through the reverse phase column and were not analyzed (Protein Analysis by Mass Spectrometry). With the high sequence coverage from mass spectrometry analysis and detection of C-terminal sequence in the anti-SEKDEL Western blot, it is confirmed that the PTH-Fc produced transiently in *N. benthamiana* is intact. It is worth noting that there is a band right below the intact PTH-Fc band on the anti-SEKDEL Western blot (Figure 4 B, lane 1), which is not observed in the anti-PTH Western blot (Figure 4 A, lane 1). Thus, the lower band corresponds to a cleaved PTH-Fc containing the C-terminal sequence (Fc domain) but not the PTH domain. This observation is consistent with a previous study on an Fc fusion protein produced in plants, where proteolytic degradation occurred within the linker domain, especially when the Fc N-glycosylation site was removed making the flexible linker domain more accessible to proteases (Xiong et al., 2019).

**Figure 4.**
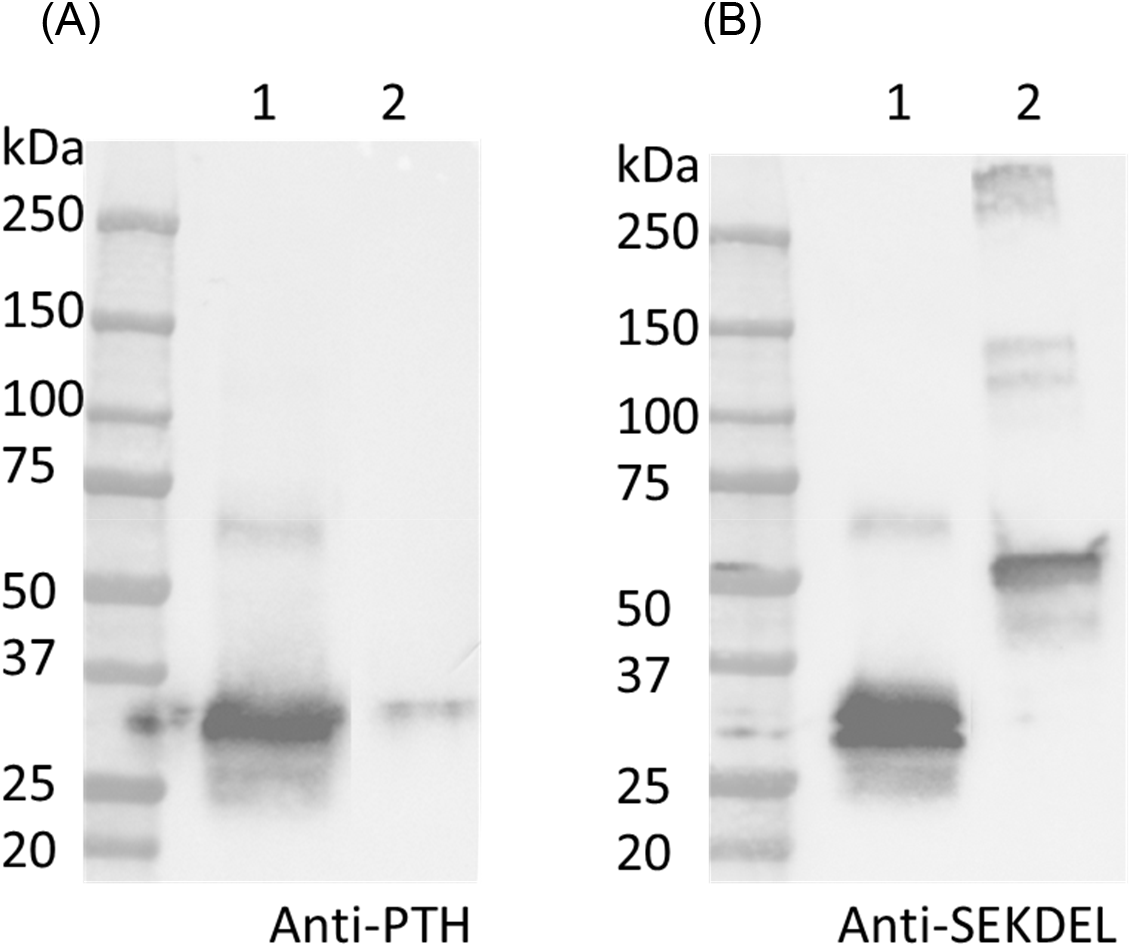
Western blot analyses detecting PTH (A) and SEKDEL (B) sequences for purified PTH-Fc (lane 1) and a control protein (lane 2, 47.2kDa) with the SEKDEL C-terminal motif. From left to right: molecular weight ladder, PTH-Fc and a control protein with SEKDEL C-terminal motif.

To further examine the protein integrity, SDS-PAGE analysis under reducing (R) and non-reducing (NR) conditions were performed on purified PTH-Fc (Figure 5). Under reducing conditions, two distinct bands between 25 – 37 kDa were observed, consistent with the anti-SEKDEL Western blot, with a band density distribution of 52% and 48% based on gel densitometry between the higher and lower bands, respectively. The concentration of PTH-Fc in the functional assays described below was based on the intact PTH-Fc only. Under non-reducing conditions, both bands dimerized, forming a wide band between 50 and 75 kDa that was not fully resolved due to their similar molecular weights. The dimerization of the lower band confirmed that protein cleavage happened within the linker domain, as protein dimerization was driven by the two cysteine residues within the Fc domain right next to the linker sequence.

**Figure 5.**
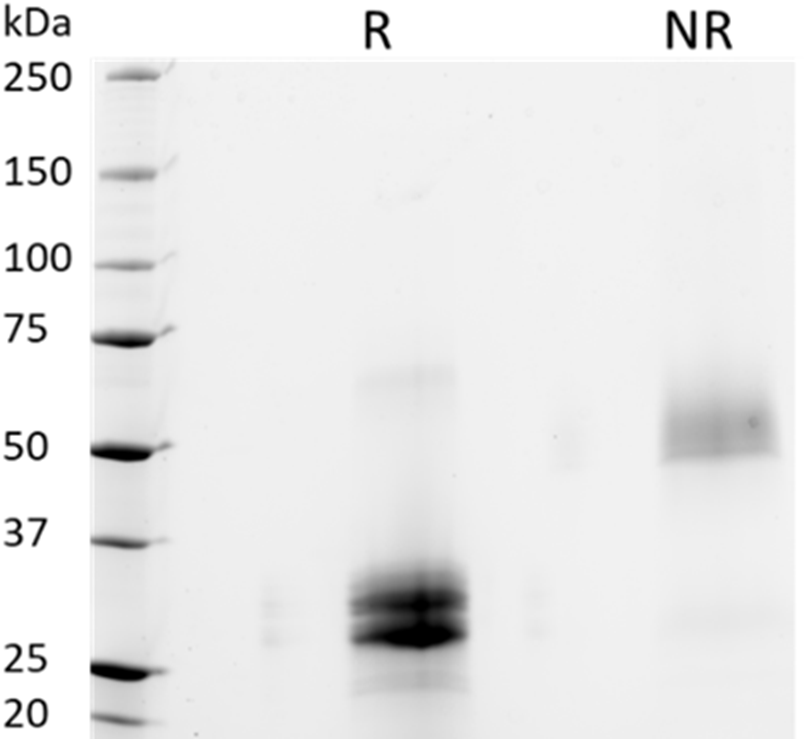
SDS-PAGE analyses of purified PTH-Fc under reducing (R) and non-reducing (NR) conditions.

### 3.4 Binding Affinity Measurement between PTH-Fc and PTH1R

Once the protein was purified and the amino acid sequence confirmed, functional assays were performed to evaluate the biological activity of PTH-Fc.

The binding between PTH-Fc and PTH1R was monitored in real-time with BLI and compared to the PTH 1-34 amino acids (positive control) and PTH1R interaction. A negative control (CMG2-Fc) was in place to rule out possible contribution from the Fc domain of PTH-Fc. The BLI sensorgrams are shown in Figure 6 A – C with the association and dissociation phases divided by the dotted read line. For both PTH-Fc and PTH trials, there was significant protein binding. For the negative control, no protein binding to the biosensor was observed, indicating the Fc domain did not non-specifically bind to PTH1R.

**Figure 6.**
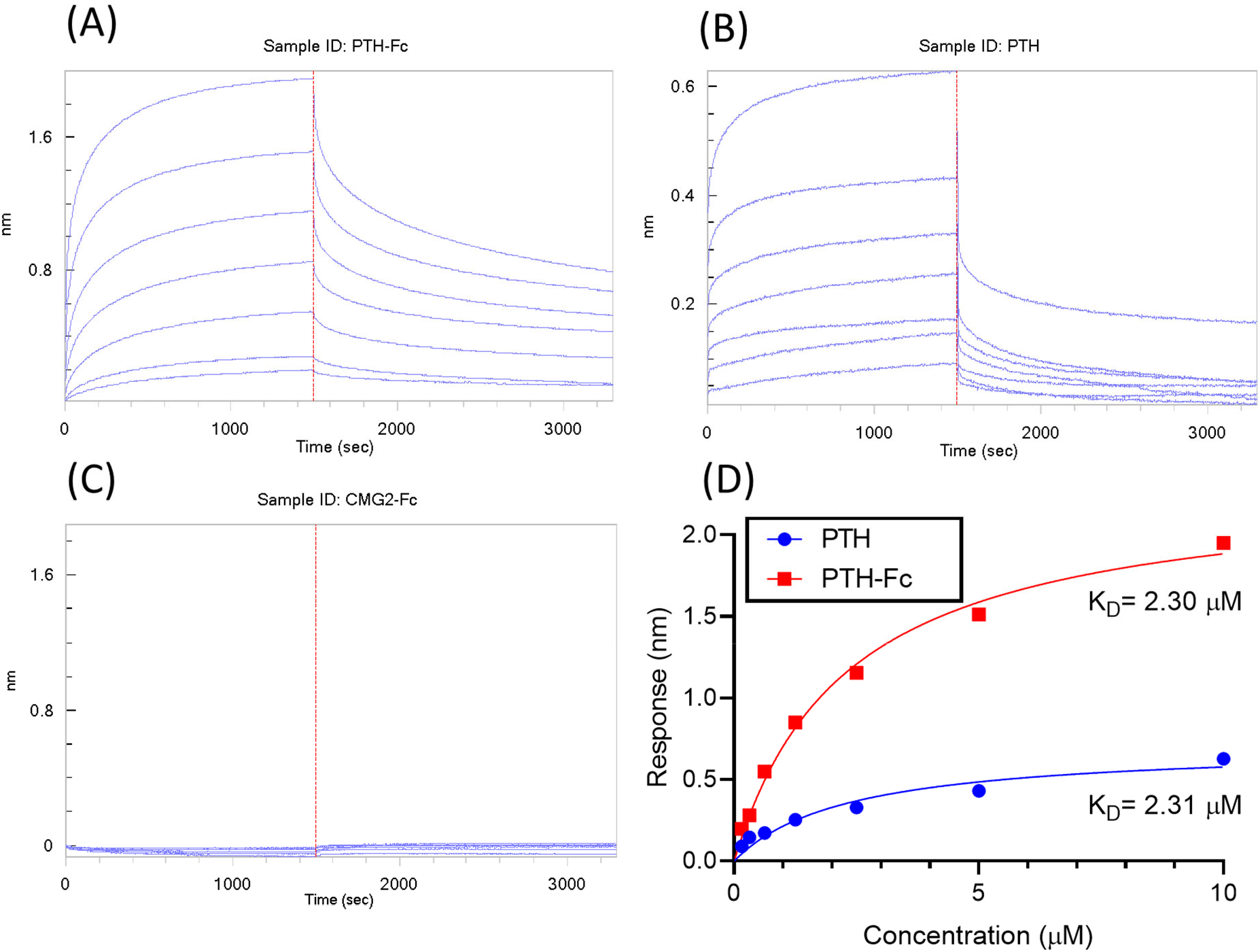
BLI sensorgrams (A) – (C) obtained from interactions between PTH1R and PTH-Fc, PTH or CMG2-Fc; (D): BLI steady state analysis of receptor binding affinity to PTH-Fc (red) and PTH (blue).

**Figure 7.**
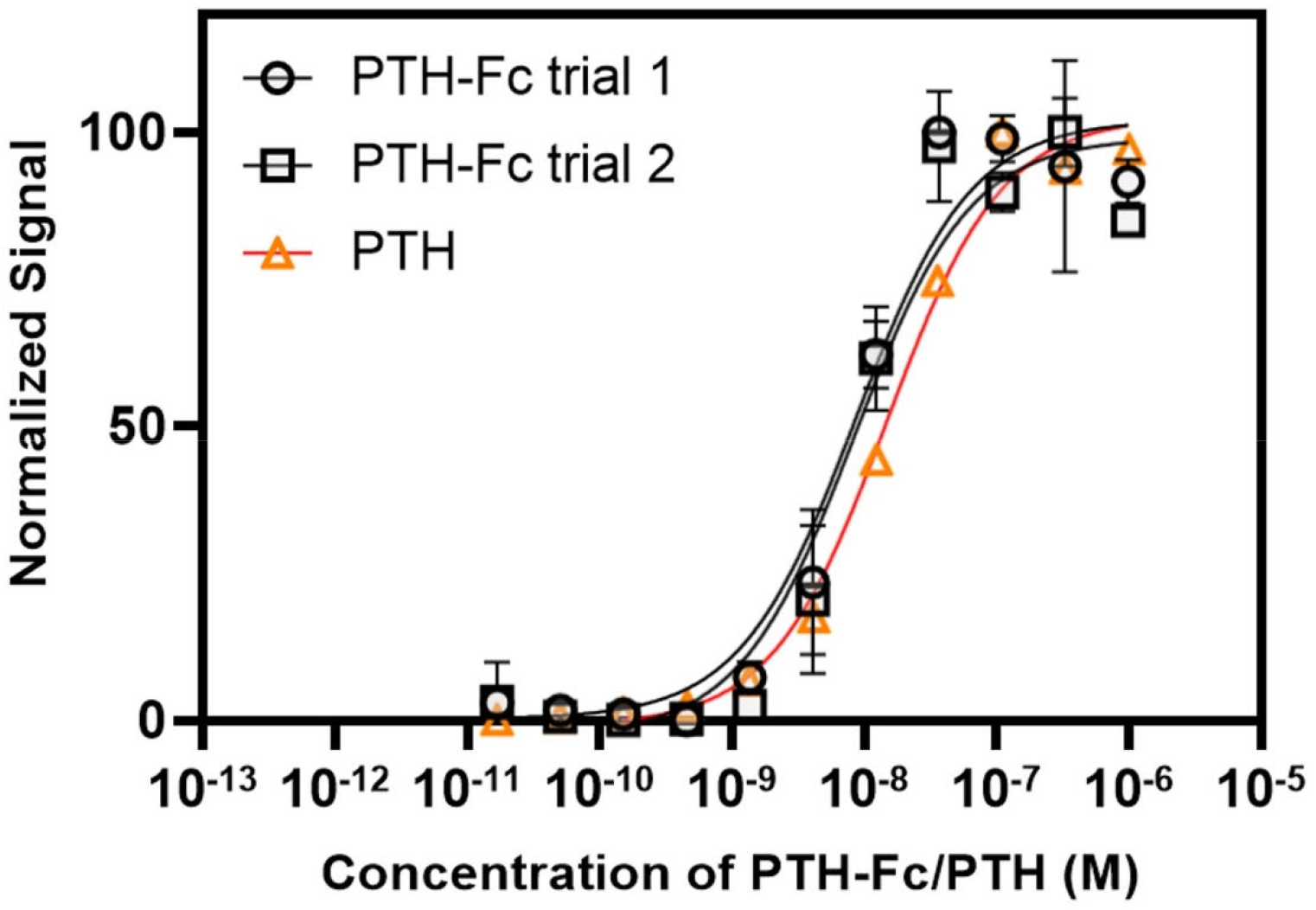
Dose-response curve from PTH-Fc or PTH treated CHO-K1 cells expressing PTH1R.

To obtain binding kinetics information, sensorgrams are usually fitted to a suitable binding kinetics model. However, in this case, as both association and dissociation happened very fast, mass transfer of protein molecules to the biolayer became the rate limiting step at the curve front, making binding kinetics fitting inaccurate. Thus, a steady state analysis was performed by plotting the average response at the end of the association phase as a function of analyte concentration, to estimate the affinity constant, K_D,_ by fitting the curves to Equation 1. The steady state analysis results are presented in Figure 6 D. At the same molar concentration, PTH-Fc elicited a higher response than PTH due to its higher molecular weight. The resulting binding affinity between PTH-Fc and PTH1R was 2.30 µM, very close to the affinity between PTH and PTH1R (2.31 µM). Those results suggest that the Fc-fusion of PTH-Fc does not interfere the binding capacity of PTH domain, and it binds to PTH1R with a similar affinity as compared to its native form, PTH. This binding affinity assay confirmed the activity of this plant recombinant PTH-Fc on protein level.

**Equation 1**. Dissociation rate constant equation. [L]: unbound ligand; [A]: unbound analyte; [LA]: ligand – analyte complex; R_max_: response when all Ls are occupied; R: response at the end of the association phase.

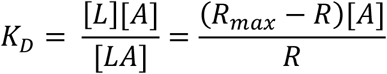

### 3.5 Receptor Stimulation Cell-based Assay

PTH anabolism is initiated by binding to PTH1R and the subsequent activation of cAMP and protein kinase A (Kostenuik et al., 2007). In this study, the cellular cAMP level was monitored within PTH-Fc treated cells as a measurement of receptor stimulation by a chemiluminescent reaction.

The dose-response curves from PTH-Fc treated cells are summarized in Figure 6.6 with highly reproducible responses from two experiments performed on different days. The EC_50_ values were estimated to be 8.62 × 10^−9^ M and 8.45 × 10^−9^ M for PTH-Fc trial 1 and trial 2, respectively. The EC_50_ from the control peptide, PTH 1-34 amino acids, was 1.49 × 10^−8^ M, calculated based on the control curve by Eurofins (PTH (1-34)). The close EC_50_ values from PTH-Fc and PTH treated cells demonstrate that the receptor stimulation potency of PTH-Fc is similar to PTH in a cell culture environment.

## 4 Discussion

In this study, PTH-Fc, an Fc-fusion of a bone regenerative protein was produced, purified and has its biological function examined in a protein-protein interaction assay and in a cell-based receptor stimulation assay. By designing PTH as an Fc-fusion, the circulatory half-life was enhanced by 33-fold (Kostenuik et al., 2007), which significantly reduces the injection frequency, making it a potential biologic for space applications. In addition, the Fc domain allows for a single step affinity purification of PTH-Fc with no native protein impurities as shown in the SDS-PAGE analysis. However, we have observed protein degradation evidenced by the double band on SDS-PAGE, which is a common issue when the protein of interest contains less structured and flexible regions (the linker) that are often targets of proteases in plant cells (Song et al., 2012). Harvesting the protein at an earlier time point might help to increase the percentage of intact protein at the cost of protein yield. For such a target, preserving the glycosylation site can reduce protein degradation by steric hinderance of oligosaccharides. Alternatively, removing or redesigning the linker sequence by avoiding proteolytic sensitive sequences or making the linker less flexible might be beneficial.

Despite the protein degradation, a significant amount of intact PTH-Fc was produced in *N. benthamiana* within 5 days. The purified protein was confirmed to behave similarly to its native form, PTH (1-34), in the PTH1R binding assay and the PTH1R stimulation CBA. The addition of the Fc domain did not interfere with its binding to PTH1R and such an approach may be applied to other biologics for half-life enhancement and simplifying the protein purification scheme, which is especially valuable under resource constraints, like in space. On the production platform side, plants are an ideal platform to produce biologics either in the long term by generating stable transgenic plants or on an on-demand basis by transiently expressing biologics via agroinfiltration or other DNA delivery methods. With its high level of flexibility and low equipment requirement, plant-based protein expression systems can contribute to making long term space missions safer and more reliable.

## 5 Summary and Future Perspectives

In this study, PTH-Fc was produced in plants transiently with an expression level of 373 ± 59 mg/kg leaf fresh weight, corresponding to an intact protein level of 192 mg/kg leaf fresh weight. The protein function was confirmed in the BLI experiment with an affinity to PTH1R of 2.30 × 10^−6^ M. In the CBA, PTH-Fc stimulated PTH1R to produce cAMP with an EC_50_ of (8.54 ± 0.12) × 10^−9^ M. In summary, this plant recombinant PTH-Fc is functional with a similar potency compared to PTH (1 – 34). Preclinical studies will help to determine the efficacy of this novel PTH *in vivo*.

To understand and prevent protein degradation, the degraded product should be N-terminal sequenced to identify the cleavage site along with testing new linkers. In addition, targeting the protein to a different subcellular location can have an impact on protein accumulation level as the protease type and level varies among cellular compartments (Benchabane et al., 2008; Pillay et al., 2014).

## 7 Author Contributions (subject to change)

YX, HH and KM designed and executed the experiments. YX wrote the initial manuscript draft. NL, KM, SN, NL edited the manuscript draft. All authors read, revised, and approved the manuscript.

## 9 Acknowledgments

We thank NASA and TRISH for the support of this study.

## 10 Data Availability Statement

The original contributions presented in this study are included in the article, further inquiries can be directed to the corresponding author.

